# P3HT-GRAPHENE DEVICE FOR THE RESTORATION OF VISUAL PROPERTIES IN A RAT MODEL OF RETINITIS PIGMENTOSA

**DOI:** 10.1101/2022.09.14.507903

**Authors:** Simona Francia, Stefano Di Marco, Mattia L. DiFrancesco, Davide V. Ferrari, Dmytro Shmal, Alessio Cavalli, Grazia Pertile, Marcella Attanasio, José Fernando Maya-Vetencourt, Giovanni Manfredi, Guglielmo Lanzani, Fabio Benfenati, Elisabetta Colombo

## Abstract

Retinal degeneration is one of the prevalent causes of blindness worldwide, for which no effective treatment has yet been identified. Inorganic photovoltaic devices have been investigated for visual restoration in advanced stage *Retinitis pigmentosa* (RP), although lack of implant flexibility and foreign-object reactions have limited their application. Organic photoactive retinal prostheses may overcome these limitations, being biomimetic and tissue friendly. Inspired by organic photovoltaic strategies involving graphene, a hybrid retinal prosthesis was recently engineered consisting of a dual poly-3-hexylthiophene (P3HT) and graphene layer onto a flexible substrate. Here, this hybrid prosthesis was subretinally implanted *in vivo* in 5-month-old Royal College of Surgeons (RCS) rats, a rodent model of RP. Implanted dystrophic rats restored visual performances at both subcortical and cortical levels in response to light stimuli, in the absence of marked inflammatory responses. Moreover, the analysis of the physical-mechanical properties after prolonged permanence in the eye showed excellent biocompatibility and robustness of the device. Overall, the results demonstrate that graphene-enhanced organic photovoltaic devices can be suitably employed for the rescue of retinal dystrophies and supports the translation of the organic strategy into the medical practice.

**TABLE OF CONTENTS:** Inspired by organic photovoltaic, a hybrid retinal prosthesis consisting of poly-3-hexylthiophene (P3HT) and graphene on a flexible substrate was subretinally implanted *in vivo* in Royal College of Surgeons (RCS) rats, a model of *Retinitis pigmentosa*. Implanted dystrophic rats restored visual performances at both subcortical and cortical levels in response to light stimuli, in the absence of marked inflammatory responses.

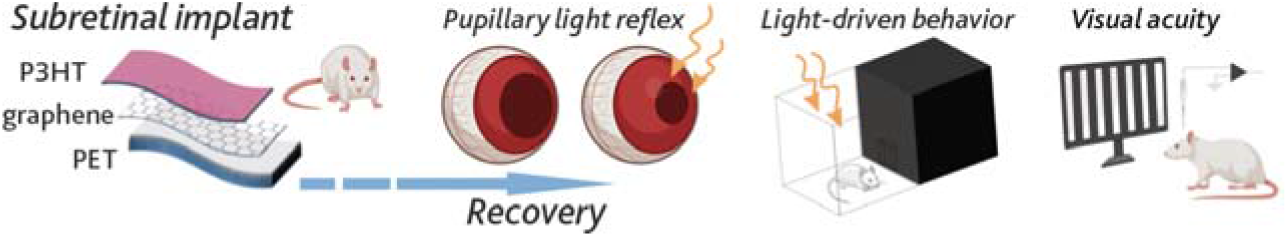

## 1. INTRODUCTION

Since its discovery, graphene has been extensively studied in connection with solar cell technology owing to the duality of its transparency coupled with extraordinary electrical properties. Graphene has been employed for different purposes over the last decade to improve organic solar cell performance, from being exploited as a photoanode ^[1]^, to a high-performance hole-transport layer for hybrid organic-inorganic solar cell systems ^[2]^, to a cathode in inverted solar cell configurations ^[3]^ or electron-transport layer ^[4]^.

Graphene-based materials interfaced with donor organic semiconductors have been demonstrated to enhance photon collection, exciton generation and charge separation ^[5,6]^. In these heterostructures, the junction between graphene and semiconducting polymers generates a built-in potential improving the transport of charge carriers and thus the extraction of photogenerated charges. The numerous attempts to enhance the phototransduction properties of organic photovoltaic devices by using graphene layers or graphene composite materials contributed to closing the gap between silicon-based and organic solar cells technologies ^[5,7]^.

On the other hand, organic photovoltaics provide high biocompatibility and mechanical compliance, paramount features for any implantable biomedical device ^[8]^. Conformability becomes a critical requirement particularly for retinal prostheses, in which the physiological curvature of the retina, together with the continuous movement of the eyeball, imposes mechanical stress and consequent risk of device failure and discomfort for the patients.

Retinal prostheses are among the best possible alternatives to the current visual rehabilitation and neuroprotective treatments aimed at rescuing vision in retinal neurodegenerative diseases. Unlike gene therapy, artificial retina devices do not rely on specific genes, and can be considered a solution in the ultimate stages of the disease, in which gene/cell replacement strategies fail or become impractical. Most retinal prosthetic technologies rely nowadays on inorganic silicon-based devices ^[9–11]^, facing problems such as need for a power supply and/or an external camera, scarce biocompatibility, poor contact with the tissue due to rigidity of the device and motion of the eye, limited number of pixels, high impedance levels and heat production ^[8]^.

To overcome these problems and inspired by organic solar cells, we recently demonstrated the potential of conjugated polymers (CPs) as planar or injectable retinal prosthetics for visual restoration in the blind, by obtaining a nearly complete recovery of visual functions in a genetic rat model of *Retinitis pigmentosa* (RP) ^[12–14]^.

To improve the performances of the organic device, we recently engineered a hybrid photosensitive interface composed by P3HT spin-coated on a chemical vapor deposition (CVD) graphene layer on polyethylene terephthalate (PET), a substrate compatible with graphene transfer technology ^[15]^. The device showed the ability to reduce firing activity *in vitro* in primary neurons and restore light sensitivity *ex vivo* in blind retinal explants. The results obtained by the prosthetic prototypes in subretinal contact with blind RCS retinal explants could reintroduce ON-like retinal ganglion cell (RGC) responses in a degenerated retina mostly ruled by OFF RGCs ^[16]^, mimicking a more physiological balance between ON/OFF-like responses. Moreover, the comparison between this prototype and the previous version, consisting of a thin film of PEDOT:PSS in place of CVD graphene, showed a clear enhancement of both charge separation and the ensuing neuronal modulation upon illumination when graphene was employed.

Here, we functionally characterize this hybrid graphene/conjugated polymer neuroprosthesis upon *in vivo* subretinal implantation in blind RCS rats. We evaluate the visual rescue provided by the enhanced device in behavioral and electrophysiological experiments, showing significant recoveries of pupillary reflex, light sensitivity and visual acuity. Post-mortem investigation of the inflammatory response in the retinal tissue contacting the device reveals no difference with respect to the levels of inflammation induced by retinal surgery and ongoing degeneration. Finally, we analyze the physical-mechanical properties of the device after permanence in the eye of the animals for 120 days to verify the robustness of the technology after long-term exposure to the biological environment.

These results demonstrate the potential of conjugated polymers and graphene in the realization of a powerful photosensitive neural interface. While the device shows excellent performance as retinal prosthesis, the application of the presented neural interface might be extended to other neurodystrophies and widen the applicability of graphene-enhanced organic photovoltaics.

## 2. RESULTS

### 2.1 Subretinal implantation of a P3HT-G device in a rat model of retinal dystrophy and its physical-mechanical characterization

We adapted a previously engineered photosensitive graphene-based device, characterized by our group *in vitro* and *ex vivo* ^[15]^, for *in vivo* subretinal implantation in RCS rats. We prepared a planar retinal device composed by a photosensitive layer of poly-3-hexylthiophene P3HT (30 mg/ml in dichlorobenzene, Sigma-Aldrich), spin-coated on a CVD graphene layer (Graphenea) transferred onto a 23 µm-thick PET substrate (P3HT-G; **Fig. 1**). PET was chosen for its long-established employment in bio-medical applications and its compatibility with CVD graphene layer transfer technology. The trapezoidal shape of the device, obtained via a laser-assisted technique, was inspired by the previous fully organic prototypes developed by our group ^[12,17]^. Both photosensitive devices (P3HT-G) and PET substrates alone (Sham) of the same size were fabricated for comparison.

**Figure 1.**
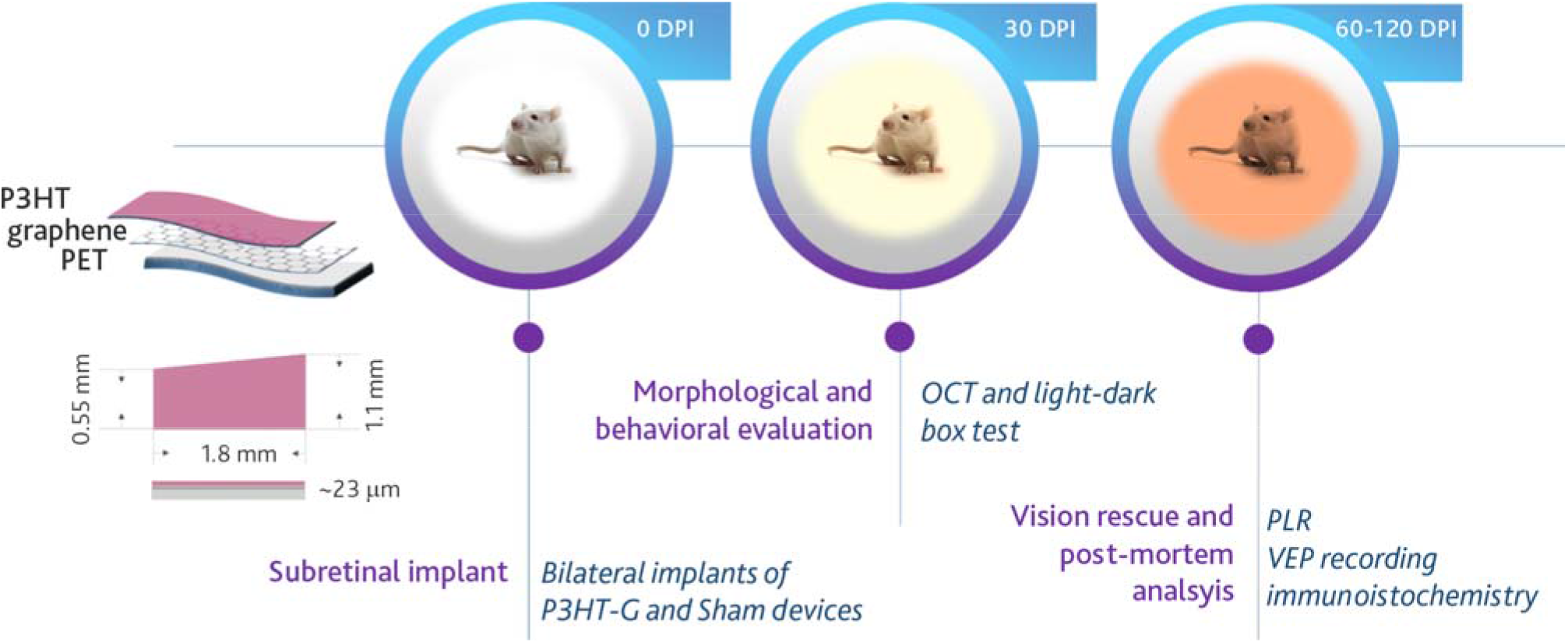
Schematic representation of the retinal prosthesis architecture and timeline of the *in vivo* experiments. Five months-old Royal College of Surgeon (RCS) rats were subretinally implanted with either P3HT-G or Sham devices. At 30 days after the implant (DPI), dystrophic RCS rats either non-implanted or implanted with P3HT-G or Sham devices, and healthy congenic rdy rats, underwent optical coherence tomography (OCT) scans and behavioral testing (light-dark box). At 60 DPI, Pupillary Light Reflex (PLR) was performed. At 120 DPI, all groups were subjected to visually evoked potential (VEP) recording in response to patterned stimuli. Thereafter, eyes were dissected for immunohistochemistry.

P3HT-G or Sham devices were bilaterally implanted in 5-month-old RCS rats, an age at which their visual functions are fully impaired ^[18,19]^. The implants were placed in the temporal retina in subretinal configuration, between the retinal pigment epithelium (RPE) and the inner nuclear layer (INL), where a proper stimulation of second-order neurons can drive visual information processing. The trans-scleral insertion of the devices in the subretinal space was possible thanks to a localized retinal detachment induced by a mild flux of viscoelastic material, in compliance with standard clinical retinal practice. At 30 days post-implant (DPI), the morphology of dystrophic RCS implanted retinas were monitored *in vivo* by optical coherence tomography (OCT) to evaluate the retina reattachment in the proximity of the implants and the integrity of retinal layers. No relevant retina damage was observed in the temporal area of the implanted animals (**Fig. 2a**, *left*) comparable to the status of non-implanted retinas, showing the safety and biocompatibility of the device. Few animals displayed a partial swelling of retinal tissue that was attributed to the physical-mechanical properties of the device (**Fig. 2a**, *right*).

**Figure 2.**
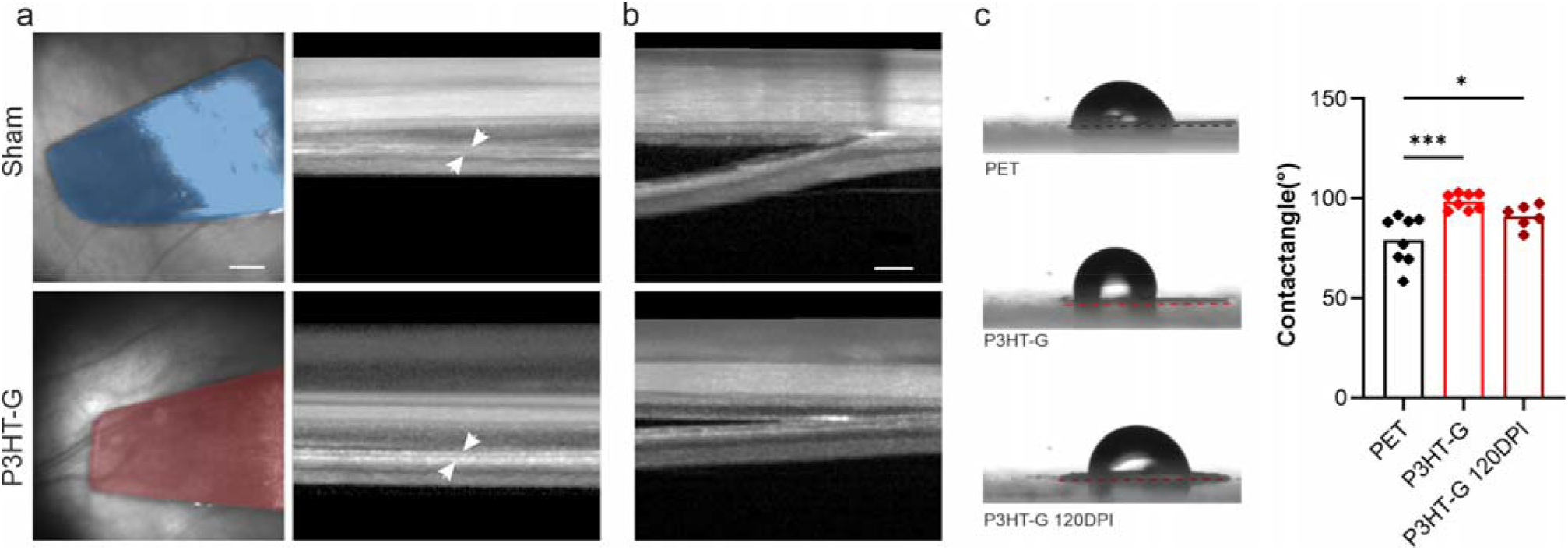
Subretinal placement of the P3HT-G device and surface characterization. **(a)** Representative *fundus* images of the retina of Sham (light blue) and P3HT-G (magenta) implanted rats showing the sagittal view of the prosthetic placement at 30 DPI (left; scale bar, 1 mm), and the respective OCT images showing the transversal section of the retinas and the devices positioned subretinally (right, white arrowheads; scale bar, 200 µm). **(b)** Representative OCT images of the few animals showing swelling of the tissue behind both Sham and P3HT-G implants (scale bar, 200 µm). **(c)** Contact angle measurements on PET and P3HT-G after fabrication and after 120 DPI (n = 8, 8 and 6 devices, respectively for PET, P3HT-G and P3HT-G/120 DPI). *p<0.05, ***p<0.001. One-way ANOVA test/Tukey’s tests. Data are shown as means ± sem with superimposed individual points.

The contact angle measurements of P3HT surface as-deposited, right after the device fabrication, showed indeed a low wettability of P3HT with respect to PET (**Fig. 2b**). This feature might have contributed to the retinal swelling observed in some animals due to a slower reabsorption of the viscoelastic material. However, most of the devices after 120 days of permanence in the eye showed a 10% improvement in wettability, attributed to the already reported oxygen-induced photoactivated doping and subsequent reduction of surface hydrophobicity ^[20]^. Therefore, although the retinal prosthesis presented some misplacement in some animals, P3HT proved to be a suitable photosensitive material for retinal applications that can easily adapt to the extracellular environment of the subretinal space.

P3HT-G devices also showed an improved adhesion on the graphene/PET substrate after 120 DPI with respect to the one obtained immediately after fabrication, as depicted in **Figure 3** by nanoindentation load-displacement measurements. The devices were glued on glass slides and subjected to a progressive load of 30 to 500 mN, while the stage was displaced transversally (**Fig. 3a**, *top*). The scratch was observed under the microscope to associate the onset of the surface damage with the corresponding critical load. As expected, a typical coating failure resulted in the delamination of the layer (**Fig. 3a**, *bottom*). P3HT-G devices after 120 DPI *in vivo* showed a significant 3-fold increase in the critical load with respect to the as-grown devices (**Fig. 3b**), confirming that the combination of CVD graphene and conjugated polymers represents a reliable strategy for the realization of a long-lasting retinal prosthesis.

**Figure 3.**
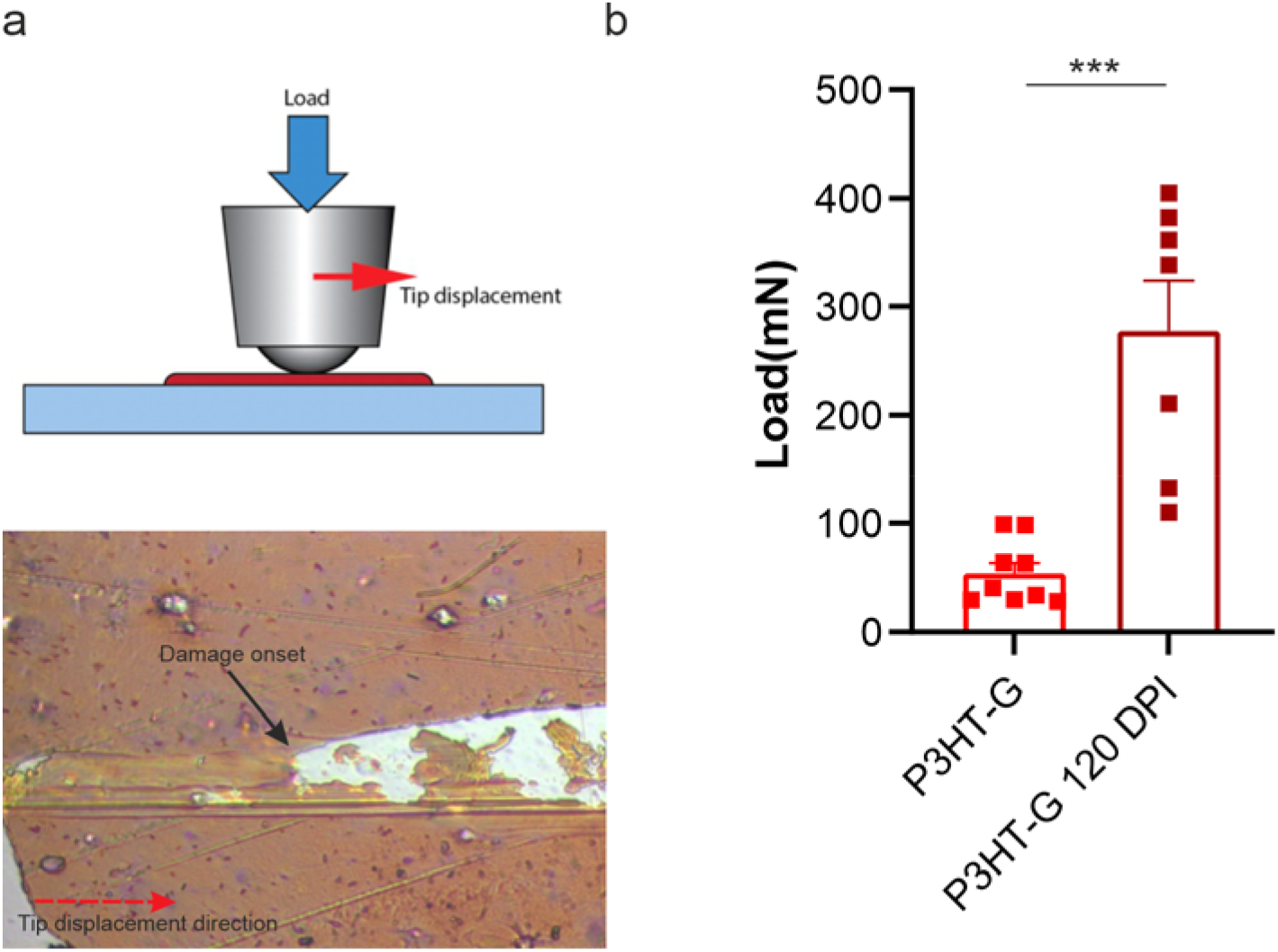
Nanoindentation experiments reveal an improved adhesion of the P3HT after permanence in the eye. **(a)** Schematic representation of the experimental setup (top), and representative image of the device surface damage after indentation (bottom). **(b)** Load quantification on graphene-enhanced retinal prostheses before implantation (P3HT-G) and after being subretinally implanted in RCS rats for 120 DPI (P3HT-G/120 DPI). Data are means ± sem with superimposed individual points. ***p<0.001, unpaired Student’s *t*-test (n= 9 and 7 for P3HT-G and P3HT-G 120 DPI respectively).

### 2.2 P3HT-G devices do not elicit a relevant inflammatory response

At 120 DPI, we investigated the extent of retinal inflammation in dystrophic RCS rats implanted with either Sham or P3HT-G devices and compared it with the retinas of aged-matched non-implanted RCS and healthy rdy controls, by analyzing markers of astrocyte/Muller gliosis (GFAP) and microgliosis (Iba-1). The analysis of GFAP immunoreactivity revealed that the P3HT-G devices did not induce further astrogliosis with respect to that triggered by photoreceptor degeneration (non-implanted and Sham-implanted RCS rats) (**Fig. 4a,b**), demonstrating that both conjugated polymers and graphene are biocompatible materials for the assembly of a retinal interface. We also observed an increased number of Iba-1 positive cells in dystrophic RCS rats compared to healthy rdy controls, regardless of the presence of the P3HT-G or Sham implant (**Fig. 4c,d**), as expected from the degeneration stage of the retina at this age. The immunohistochemical characterization demonstrated that the partial lack of reabsorption of the liquids generated during surgery in some animals at 30 DPI did not induce any inflammatory reaction and possibly recovered to a physiological condition, also favored by the device adaptation to the retinal environment.

**Figure 4.**
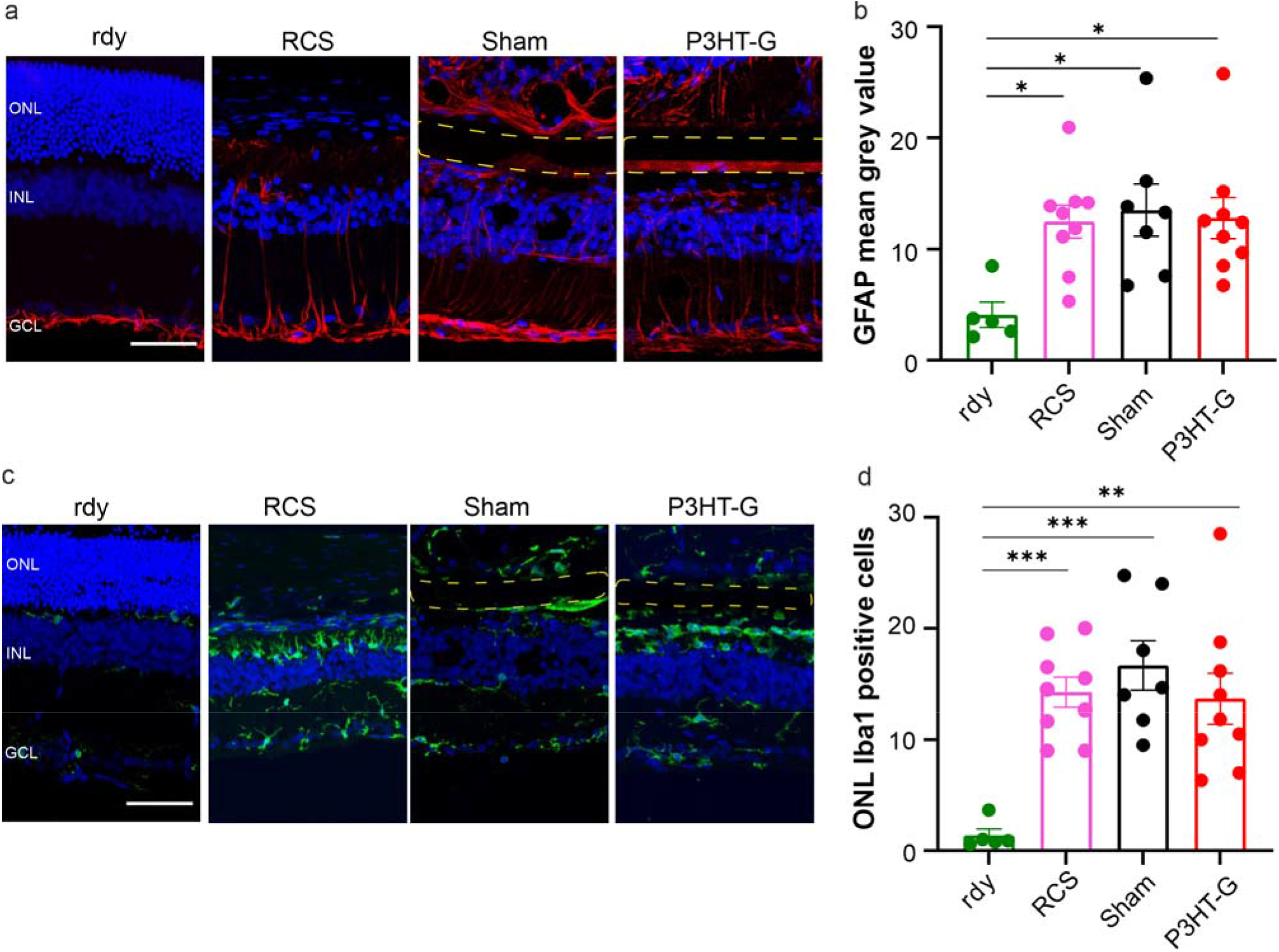
P3HT-G prostheses do not elicit retinal inflammatory responses in RCS rats. Representative transversal sections of retinas dissected from non-dystrophic controls (rdy) and from dystrophic RCS rats that were non-implanted or implanted with either Sham or P3HT-G device. **(a)** Representative retinal sections immunolabelled for the astrocyte/Müller cell marker GFAP (red) merged with bisbenzimide nuclear labelling (blue). The dashed lines indicate the placement of the devices (scale bar, 50 µm). Note the P3HT autofluorescence of the P3HT-G device. **(b)** Quantitative analysis of GFAP immunoreactivity in the inner retina (mean fluorescence intensity). **(c)** Representative retinal sections immunolabelled for Iba-1 (green) merged with bisbenzimide nuclear labelling (blue). The dashed lines indicate the placement of the devices (scale bar, 50 µm). **(d)** The histogram shows the number of Iba-1 positive cells in the ONL from at least two fields per temporal retina section. No differences were detected between the immunoreactivities of temporal and nasal sections. Data are means ± sem with superimposed individual points. *p<0.05, **p<0.01, ***p<0.01, one-way ANOVA/Tukey’s tests (sample size: n= 5, 9, 7, 9 for rdy, RCS, Sham, and P3HT-G, respectively).

These results, in line with previous works ^[12,21]^, confirm that the implantation of our retinal device is associated with a full retinal recovery from the surgical stress, with no additional inflammation caused by the presence of graphene.

### 2.3 RCS rats implanted with the P3HT-G device present visual rescue at subcortical and cortical levels

Next, we investigated the extent of visual restoration by the P3HT-G retinal prosthesis at 60 DPI by analyzing the Pupillary Light Reflex (PLR), an experiment measuring the constriction and subsequent dilation of the pupil in response to light stimuli ^[22]^. We analyzed PLR dynamics in RCS rats implanted with Sham and P3HT-G devices and compared them to the responses of aged-matched healthy rdy and non-implanted dystrophic RCS rats. The anesthetized animals were subjected to monocular flashes of green light (see Methods), while videorecording pupil dynamics with an infrared camera (**Fig. 5a**, *left*). We evaluated subcortical responses to light stimulation in the four experimental groups by measuring PLR latency, constriction extent and rate, as well as post-stimulus pupillary dilation (PIPR) at 5, 20 and 50 lux (**Fig. 5a**, *right*).

**Figure 5.**
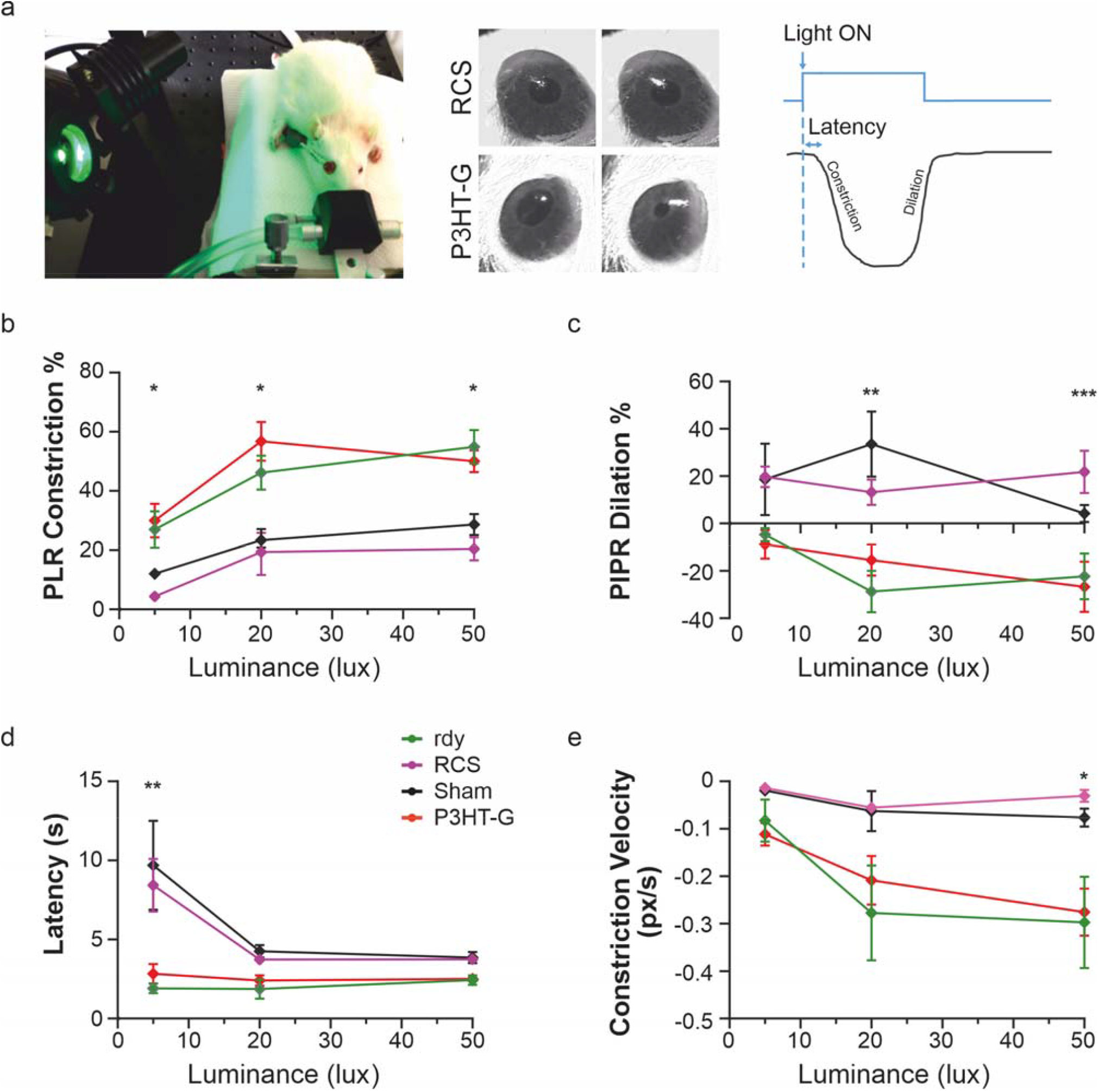
P3HT-G devices restore subcortical reflexes in RCS rats. **(a)** *Left*: Pupillary Light Reflex (PLR) setup. *Middle:* Representative images of the pupil’s maximal constriction from video-recordings performed under infrared illumination in response to 20-sec green light stimuli (530 nm; 20 lux) in RCS rats either non-implanted or implanted with the P3HT-G device. *Right:* Schematic representation of the pupillary reflex dynamics. All experimental groups were tested at 60 DPI and were challenged with light stimuli at 5, 20, and 50 lux for the percentage of pupillary constriction **(b)**, constriction velocity **(c)**, latency **(d)**, and percentage of PIPR dilation **(e)**. P3HT-G implanted RCS rats showed a significantly more intense and fast pupillary constriction, a smaller relaxation, and a shorter constriction latency than non-implanted RCS or Sham-implanted rats. Data are means ± sem. *p<0.05, **p<0.01, ***p<0.001, Kruskal-Wallis/Dunn’s tests (PLR constriction and PIPR dilation, n = 6, 6, 6, 3; PLR Latency, n = 3, 3, 3, 6; Constriction velocity, n = 3, 3, 6, 3, for rdy rats and non-implanted, P3HT-G-implanted and Sham-implanted RCS rats, respectively).

Dystrophic RCS rats subretinally implanted with P3HT-G devices showed a significantly larger pupillary constriction than blind non-implanted and Sham-implanted control groups at all luminance levels, indicating the light-evoked stimulation of inner retinal neurons induced by the graphene-based prosthesis (**Fig. 5b**). Moreover, the significantly smaller relaxation of the pupil in the animals implanted with P3HT-G devices at 20 and 50 lux indicates that the involvement of intrinsically sensitive retinal ganglion cells (ipRGCs) was also rescued to the level of healthy rdy controls (**Fig. 5c**). At 5 and 50 lux, P3HT-G-implanted RCS rats significantly recovered latency and constriction velocity, respectively, becoming indistinguishable from age-matched healthy rdy controls (**Fig. 5d,e**). To assess the restoration of visual functions and spatial discrimination at the cortical level, we recorded pattern visually evoked potentials (pVEPs) in the binocular portion of visual cortex V1 (OC1b), by subjecting the animals to switching gratings (1 Hz) of progressively higher spatial frequencies, covering a range of 0.1 to 1 c/deg (**Fig. 6a**).

**Figure 6.**
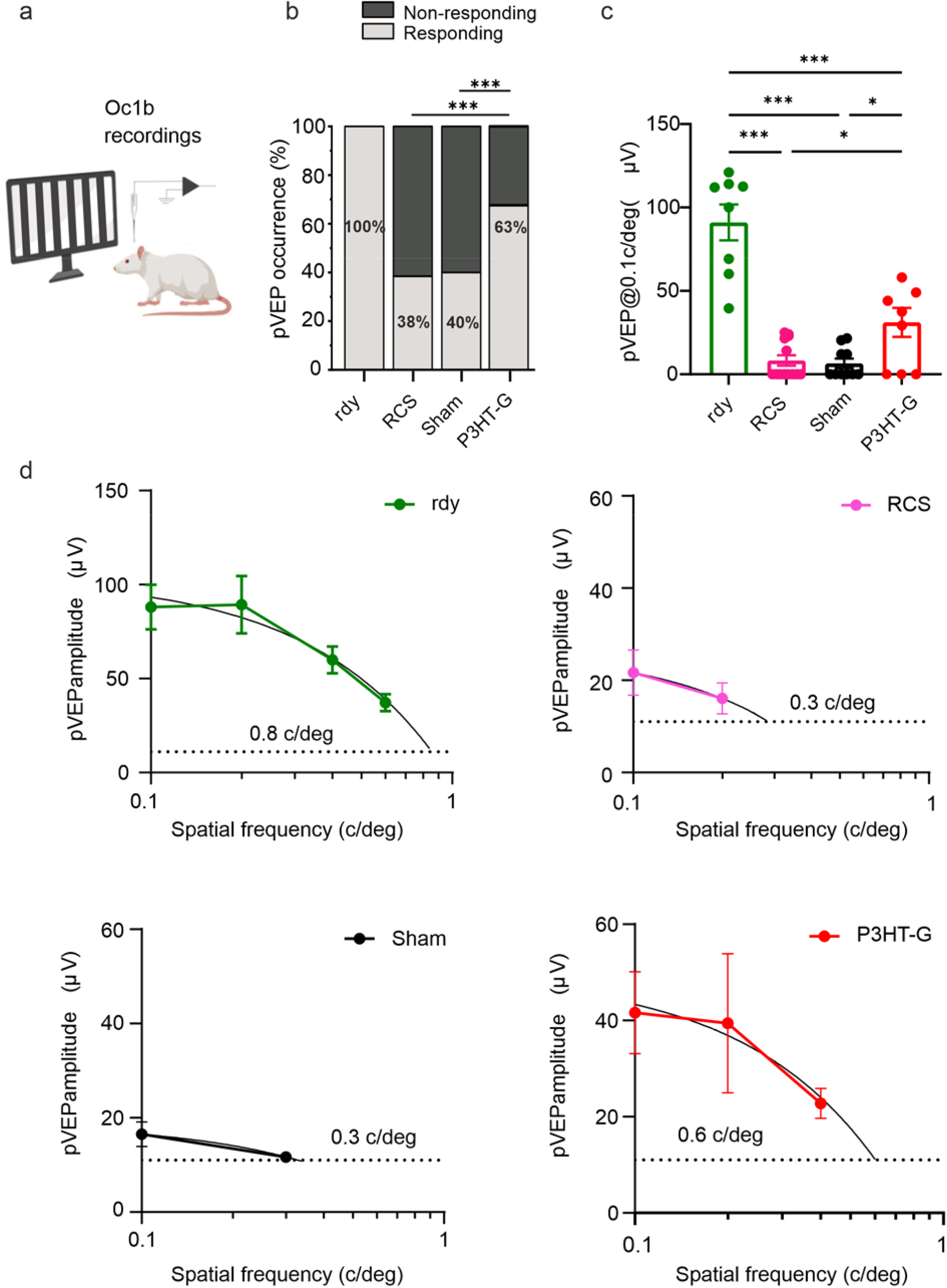
P3HT-G prosthesis restores visually evoked potentials in dystrophic subretinally implanted RCS rats. **(a)** Schematic representation of the pattern visual evoked potentials (pVEP) recording setup. The four experimental groups (healthy rdy rats, non-implanted, P3HT-G implanted and Sham-implanted dystrophic RCS rats) were recorded in the binocular portion of visual cortex V1 (Oc1b) at 120 DPI. **(b)** Percentage of animals with a pVEP amplitude above 2X the standard deviation (SD) of the noise in all groups. ***p<0.001. Fisher’s exact test (sample size: 8, 13, 10, 8 for rdy, RCS, Sham, and P3HT-G, respectively). **(c)** pVEP amplitudes in response to patterned stimuli at 0.1 c/deg showed a recovery in RCS rats implanted with P3HT-G compared to non-implanted or Sham-implanted rats. Data are means ± sem with superimposed individual points. *p<0.05, ***p<0.001, one-way ANOVA/Fisher’s tests. **(d)** Mean (± sem) pVEP amplitudes calculated for the animals with cortical responses greater than 2x SD of the noise in each experimental group. Linear regression was computed on the decreasing pVEP amplitudes before plateauing, and visual acuity was estimated from the intercept with y= 2x SD noise line (dashed line). Sample size: 8, 5, 4, 5 for rdy, RCS, Sham, and P3HT-G, respectively.

The pVEP amplitude of dystrophic RCS rats either non-implanted or Sham-implanted showed negligible responses to patterned stimuli (110 cd/m^2^, 1 Hz), testifying the severe degree of degeneration of the RP model. About 60% of the RCS rats implanted with P3HT-G devices presented a pVEP response at 0.1 c/deg, as compared to the ∼40% of non-implanted or Sham-implanted RCS rats, confirming the efficiency of the prosthetic strategy (**Fig. 6b**). Interestingly, dystrophic RCS rats implanted with the P3HT-G device improved their spatial acuity (**Fig. 6c-d**), showing a significant recovery of the cortical responses evoked by patterned stimuli at 0.1 c/deg compared to Sham-implanted or non-implanted RCS rats that showed minimal pVEP amplitudes. The recovery in VEP amplitude and spatial discrimination in RCS rats implanted with the P3HT-G device was not however a *restitutio ad integrum*, as their responses were still lower than those of age-matched healthy rdy controls (**Fig. 6c,d**). However, the recovery of visual responses at the cortical level in RCS rats highlight the efficacy of the P3HT-G device for visual restoration in RP.

### 2.4 Subretinal P3HT-G prosthesis restores light-aversion behavior in RCS rats

Finally, we evaluated the degree of visual rescue triggered by the P3HT-G device by analyzing the visually driven behavior of the animals at 30 DPI using the light-dark box test. The behavioral setup employed to evaluate the rodent innate aversion to bright open spaces is composed of two communicating “light” and “dark” chambers (**Fig. 7a**). A short escape latency from the illuminated area and a predominant permanence in the dark compartment (dark preference) are reliable indexes of physiological light perception. A significant recovery of both latency and dark preference was observed in RCS rats implanted with P3HT-G devices with respect to the Sham-implanted or non-implanted RCS rats (**Fig. 7b,c**), demonstrating the presence of the graphene-mediated phototransduction predicted *in vitro* and *ex vivo* in our previous work ^[15]^. The total number of transitions between the two compartments of the box did not show any difference across the experimental groups, testifying the absence of motor deficits that could have biased the test (**Fig. 7d**). Overall, the behavioral data confirm the conclusion that the graphene-enhanced retinal prosthesis rescues light sensitivity in the RCS rat model of RP.

**Figure 7.**
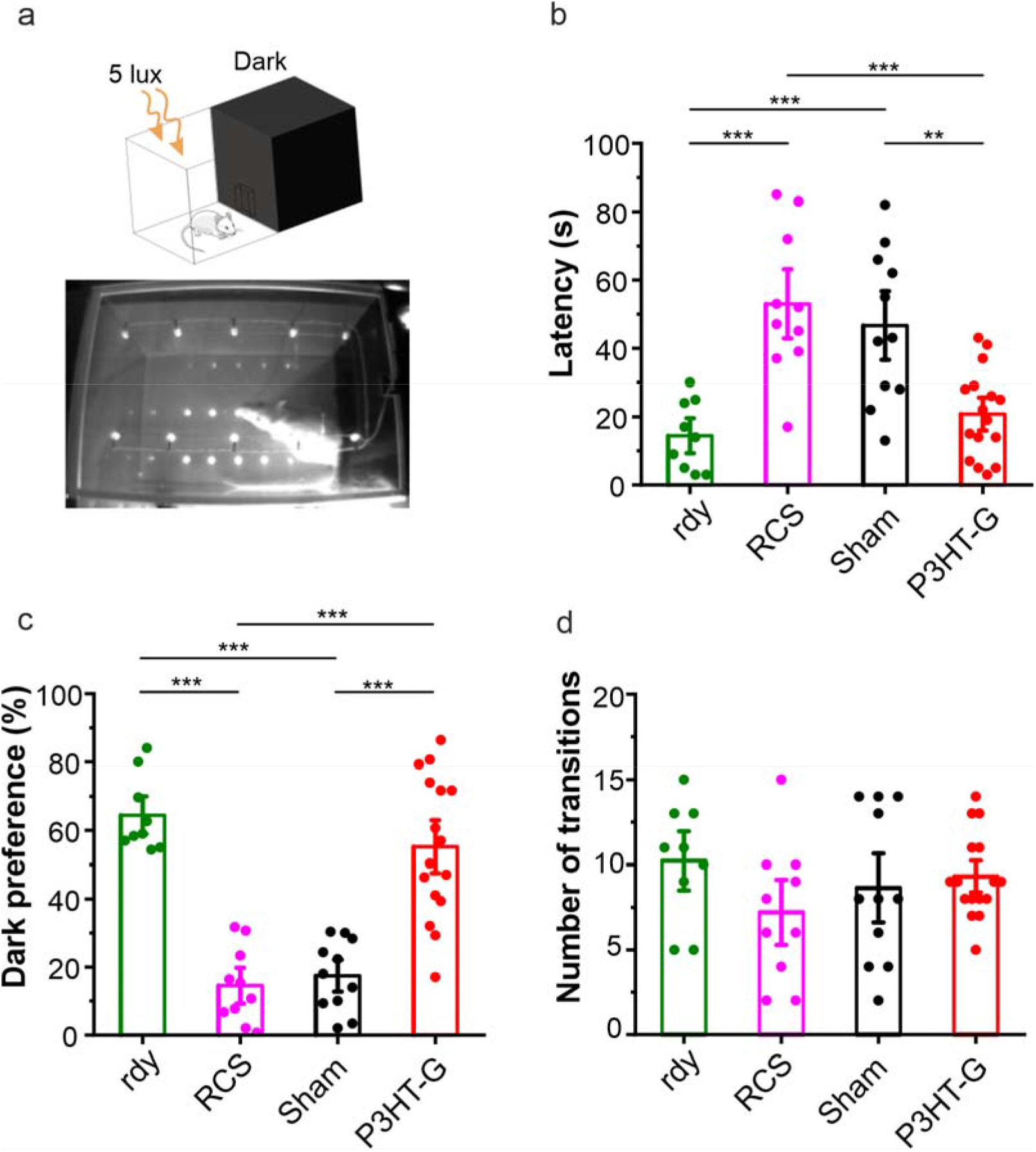
Subretinal P3HT-G prosthesis restores visually driven behavior in RCS rats. **(a)** *Top:* Schematic representation of a light-dark box setup. *Bottom:* Representative stop-frame of videorecording of a light dark box experiment. All experimental groups were analyzed at 30 DPI. **(b)** Escape time of the rats from the illuminated area. **(c)** Percent time spent in the dark compartment. **(d)** Number of transitions between the two compartments reveals the absence of motor impairments. RCS rats subretinally implanted with P3HT-G show a significant reduction of light-escape-latency, and an increased dark preference with respect to Sham-implanted or non-implanted blind RCS rats. Data are means ± sem with superimposed individual points. **p<0.01, ***p<0.001, one-way ANOVA/Tukey’s tests. (Sample size: 9, 10, 11, 16 for rdy, RCS, Sham and P3HT-G, respectively).

## 3. DISCUSSION AND CONCLUSION

Conjugated polymers have been thoroughly studied as neuronal interfaces for photostimulation *in vitro* and the rescue of visual functions in models of retina degeneration *in vivo* ^[12–14,21,23]^. Based on this premise and considering the increasing demand for high-sensitivity retinal prosthetics, we engineered a prototype of planar organic subretinal implant whose efficiency is boosted by the presence of graphene.

One of the focuses of conjugated polymer photovoltaics has been in fact the choice of the appropriate architecture of the donor-acceptor pair for an efficient excitonic dissociation upon illumination. Among the bulk heterojunction structures, CVD graphene, and more generally 2D materials, have played an important role in increasing organic solar cell efficiencies that are progressively approaching those of their inorganic counterparts ^[5,7]^.

On the other hand, the intrinsic biocompatibility of graphene, combined with the enhanced optical and electrical properties, make graphene a suitable candidate for the realization of neural interfaces for the modulation and recording of neuronal activity ^[24,25]^. Although further efforts are needed to establish hybrid 2D-on-3D heterojunction photovoltaic systems in terms of long-term durability, the mechanical compliance associated with an improved robustness of these architectures inspired their application in the bio-medical field. We recently engineered a fully organic photosensitive device composed of P3HT on CVD graphene supported by PET that effectively mediated photostimulation of neurons cultured up to 15 days *in vitro* and triggered the light-responses of RGCs in blind RCS retinal explants ^[15]^.

Here, we demonstrate the suitability of the P3HT-G prosthetic prototype to restore light sensitivity and visual acuity in a model of retinal degeneration *in vivo*. The retinal devices have been designed to fit the requirements for subretinal implantation in terms of shape and thickness, while the presence of a P3HT/graphene interface ensured the proper charge separation and consequent inner retina stimulation after light exposure.

The retinal prosthesis was implanted with the P3HT layer in contact with inner retinal neurons in 5-months-old RCS rats, an age at which the degeneration of the photoreceptors is already complete, and no residual visual functions are present. OCT scans revealed the correct subretinal placement of the devices, although revealing in some animals a mild residual swelling behind the devices at 30 DPI, attributed to a potential effect of the hydrophobicity of the implants in the first phase of tissue recovery and adaptation after surgery. However, the analysis of the contact angle of the devices after 120 DPI permanence in the eye showed a gain in the surface wettability, in line with previous characterizations of P3HT films in contact with aqueous solutions ^[20]^. Moreover, the P3HT film on graphene showed an improved scratch resistance after 120 DPI. The robustness to delamination of the device could be attributed to the concomitant effect of an improvement of the load resistance of CVD graphene on PET ^[26]^, together with the stronger π-π interaction between the P3HT backbone chain and the graphene surface ^[27]^. While a more conformable or even biodegradable substrate could be preferable, PET represents one of the best choices for CVD-graphene transfer on a flexible and organic substrate. Moreover, the tolerability of the device upon prolonged permanence in contact with retinal tissues is enhanced by the progressive decrease of the P3HT surface hydrophobicity observed in post-mortem samples.

Our study demonstrates the overall biocompatibility and reliability of the prosthetic architecture up to 120 DPI *in vivo*, supported by the substantial absence of inflammatory reactions to the presence of the implant by astrocytes and microglia. The visual function of the animals implanted with the P3HT-G devices revealed a satisfactory rescue of the pupillary light reflex dynamics in a wide range of luminance levels. In addition, cortical responses elicited in the presence of the P3HT-G prosthetics upon pattern light stimulation testify a partial recovery of visual acuity. These results, together with the restoration of light-driven behaviors, demonstrate the in vivo efficacy of conjugated polymer/graphene hybrid neuroprosthetics. The biocompatibility and long-term stability of the device *in vivo*, together with the prolonged rescue of visual functions up to four months after implantation, prove the potential of the presented strategy for the recovery of blindness and extends its application to the broader field of neurodegenerative diseases.

## Acknowledgments

This project has received funding from the European Union’s Horizon 2020 research and innovation programme under grant agreements No. 785219 (Graphene Flaghship - Core 2) and No. 881603 (Graphene Flaghship - Core 3).

## 4. METHODS

### 4.1 Ethical approval of animal manipulation and procedures

All manipulations and procedures involving animals were carried out in compliance with the guidelines established by the European Community Council (Directive 2014/26/EU of 4 March Sep 2014) and were approved by the Italian Ministry of Health (Authorization # 357/2019-PR). RCS and rdy rats were kindly provided by dr. M.M. La Vail (Beckman Vision Center, University of California San Francisco, CA). The animals were kept on a 12/12h light-dark cycle in standard breeding conditions. Experimental groups were randomly selected maintaining a balance of females and males.

### 4.2 Devices fabrication

A graphene layer, produced by Chemical Vapor Deposition (CVD) (Graphenea) was transferred on a PolyEthylene Terephthalate (PET) substrate (23 µm thickness) and cut in 2×2 cm samples. Subsequently, poly-3-hexylthiophene (P3HT, Sigma Aldrich, 15,000-45,000 molecular weight) dissolved in 1,2-Dichlorobenzene (Sigma-Aldrich) at 30 mg/ml was spin-coated on the PET surfaces by a two-step process: 800 rpm for 5 s followed by 1,600 rpm for 120 s, and annealed at 120 °C for 20 min ^[15]^. The devices were then cut to the final trapezoidal shape by a laser-assisted technique and subjected to ethylene oxide sterilization.

### 4.3 Physical-mechanical characterization

Contact angle measurements were performed with a OCAH-200 DataPhysics system. Static water contact angle (WCA) was measured at room temperature upon delivery of milli-Q water droplets of 0.2-0.5 μl through a gas-tight Hamilton precision syringe. The samples were glued to a coverslip and positioned on the testing surface placed in a horizontal position.

The coating adhesion was evaluated by scratch tests on a MicroCombi Indenter/scratch apparatus by Anton Paar, equipped with a spherical tip of 1 mm radius. After gluing the devices on glass slides, an initial load of 30 mN was applied and increased progressively to 500 mN, producing a scratch length of about 1 mm (1 mm/min). After each test, the scratch was observed under the built-in microscope to identify damage and to associate the damage onset with the corresponding critical load.

### 4.4 Subretinal implant procedures

*Anesthesia:* 5–month-old RCS rats were anesthetized with diazepam (Ziapam, Ecuphar) (5 mg/kg) by an intraperitoneal injection followed by intramuscular administration of xylazine (Rompun, Bayer) (5 mg/kg) and ketamine (Lobotor, Acme) (50 mg/kg). Then, 1% tropicamide eye drops (VISUfarma) were administered allowing a complete pupil dilation. *Device implantation*: after fixation of the eyelid, a 2 mm cut of the conjunctiva in the temporal eye allowed a *limbus*-parallel 1 mm incision through the sclera and choroid. The retina was gently detached through the incision using either hyaluronic acid or surgical scissors, and the devices were inserted in subretinal position. The scleral incision was sutured by thermally assisted coagulation. The P3HT-G device was positioned with the polymeric coating facing the inner retina. During the whole surgery, a proper hydration of the cornea and tissues was maintained. All the surgical procedures were carried out under sterile conditions using a Leica ophthalmic surgical microscope. At the end of the surgery, the status and integrity of the implanted retina was monitored using indirect ophthalmoscopy. Antibiotic and corticosteroid eye-drops (TobraDex 0.3% + 0.1%, Alcon) were applied to prevent post-surgery inflammation and/or infection. Both P3HT-G and Sham devices were implanted binocularly.

### 4.5 Optical Coherence Tomography

Intraperitoneal injection of diazepam (5 mg/kg) followed by intramuscular administration of xylazine (5 mg/kg) and ketamine (50 mg/kg) was carried out in all experimental groups. Optical coherence tomography (OCT) of the implanted and non-implanted retinas was performed using a Spectralis™ HRA/OCT device. Each two-dimensional B-Scan recorded at 30° scan angle consisted of 1536 A-Scans and an average of 100 frames. Imaging was performed using the proprietary software package Eye Explorer (version 3.2.1.0). Implanted RCS rats were subjected to the behavioral test at 30 DPI.

### 4.6 Retina immunohistochemistry

Euthanasia was carried out by CO_2_ inhalation followed by cervical dislocation. Eyes were enucleated keeping track of the eye orientation during processing. Eyes were fixed in 4% paraformaldehyde (Sigma-Aldrich) in 0.1 M phosphate-buffered saline (PBS, Sigma-Aldrich) for 6 hours, extensively washed in 0.1 M PBS, and cryoprotected by equilibration with 30% sucrose. Eyecups were obtained by removing the cornea, the iris, and the lens and then embedded in OCT freezing medium (Tissue-Tek; Qiagen), frozen in dry ice and cryo-sectioned at 25 μm using an MC5050 cryostat (Histo-Line Laboratories). Eyes were oriented by maintaining the position of the device (either Sham or P3HT-G) implanted in the temporal area of the retina. Sections were mounted on gelatin-coated glass slides and maintained at -20 °C until processed. Retina sections were incubated with 10 % normal goat serum (NGS, Sigma–Aldrich) at room temperature for 2 hr, then incubated overnight at 4 °C with the primary antibody of interest: ionized calcium-binding adaptor molecule 1 (Iba-1, 1:1000; Wako Pure Chemical Industries, Japan) and glial fibrillary acidic protein (GFAP, 1:200; Sigma-Aldrich) in 1% NGS and 0.3% Triton-X-100. The secondary antibodies were Alexa Fluor 488 and Alexa Fluor 564 (1:200; Molecular Probes, Invitrogen, Carlsbad, CA), together with bisbenzimide nuclear dye 33342 (1 μM; Hoechst), in 5% NGS and 0.3% Triton-X-100 for 1,5 hrs at room temperature. The preparations were dried and mounted with Mowiol (Sigma – Aldrich) prior to confocal imaging acquisition. Retinal sections were imaged with a Leica SP8 confocal microscope (Wetzlar, Germany).

The analyses were performed (ImageJ, NIH) by averaging measurements from at least two regions of interest (ROIs) for each slice of temporal retina. GFAP mean grey value was obtained from the fluorescence intensity sum over the central 15 slices of a z-stack acquisition for each field, comprising all retina layers. The presented Iba-1 cell count number, obtained from the z-max projection of the z-stack acquisitions, refers only to the ONL layer.

### 4.7 Pupillary Light Reflex

RCS and rdy rats were dark-adapted for 30 min, anesthetized with isoflurane (Isovet, 3% induction, 2% maintenance in oxygen) and positioned with the recorded eye exposed to infrared light (LED@780 nm, Thorlabs) for visualization of the pupil through a recording camera (Moticam 1080HDMI camera). Light stimulation from a 530 nm LED (Thorlabs) 20 s of green light exposure at 5, 20, and 50 lux, followed by 48 s recording in the darkness. The following parameters were analyzed: i) *baseline*: pupil area before the starting of the stimulus; ii) *latency*: the interval of time (msec) between the onset of the stimulus and the start of the constriction; iii) *PLR constriction:* the averaged area of the pupil during the maximal constriction normalized to the baseline area; iv) *PIPR dilation:* the averaged area of the pupil during the maximal dilation normalized to the baseline, following the stimulus offset; v) *constriction:* calculated as the minimum and maximum value of the first derivative calculated during the constrictor or dilatory phase, respectively ^[28,29]^. We discarded the pupil area measurements whose videos were not of high quality.

### 4.8 *In vivo* electrophysiology

RCS and rdy controls rats were anaesthetized with isoflurane (Isovet) (3% induction, 2% maintenance in oxygen) and placed in a stereotaxic frame (Narishige), while body temperature was monitored. The skull was perforated in correspondence with the binocular portion of V1 (OC1b), then the dura mater was removed exposing the brain surface. A glass micropipette electrode from extracellular recordings (2 – 4 MΩ) filled with NaCl 3 M, was placed in the OC1b, 4.8 - 5 mm from λ (intersection between the sagittal and lambdoid sutures). Being both eyes maintained open and fixed, they were continuously hydrated with saline solution (NaCl 0.9%) during surgery and recordings. The spatial vision and pattern perception was evaluated by recording the pattern visually evoked potentials (pVEPs; recorded at about 400 µm depths in the visual cortex), in response to patterned stimuli of 100 ms at 1 Hz in front of the animals (binocular exposure). Patterned visual stimuli consisted of horizontal sinusoidal contrast-reversing gratings of increasing spatial frequencies (0.1 to 1 c/deg). The visual stimulation was generated at a luminance of 110 cd/m^2^ (SpectroCAL MKII Spectroradiometer) by a ViSaGe MKII Stimulus Generator (Cambridge Research Systems) connected to a monitor (20×22” area, 100 % brightness and contrast) located at 25 cm from the eyes of the experimental rats. All electrophysiological signals were amplified and band-pass filtered (0.1-100 Hz) by a NeuroLog system (Digitimer), digitized through a National Instruments multifunction board (NI USB-6251), and acquired using MATLAB R2019b. pVEPs amplitudes were extracted from 200-500 sweeps average traces. A relevant VEP signal was considered whenever above 2-fold the standard deviation (SD) of the noise (peak-to-baseline amplitude). Visual acuity was estimated as the intercept with y=2x SD of the linear regression of the pVEPs amplitude curve as a log-function of the spatial frequency.

### 4.9 Light-dark box test

The behavioral trials were performed in an apparatus consisting of a “light” and “dark” compartments communicating through a small door, with the size of the illuminated area about twice that of the dark area. All experimental groups were dark-adapted for 30 min and then introduced in the “light” area with the light off. For 5 min, a video recording was performed while illuminating the “light” compartment with a 5-lux intensity to monitor: i) the latency of escape from the illuminated area, ii) the percentage of time spent in both areas, and iii) the total number of transitions the rats performed between the two compartments (monitored as an index of light-independent motor activity). Implanted RCS rats were subjected to the behavioral test at 30 DPI.

### 4.10 Statistical analysis

The sample size needed for the planned experiments (n) was predetermined using the G*Power software for ANOVA test with four experimental groups, considering an effect size = 0.25-0.40 with α = 0.05 and β = 0.9, based on similar experiments and preliminary data. Experimental data are expressed as means ± sem with individual experimental points (n). Normality tests were performed to evaluate the normal distribution of data populations and consequently parametric or non-parametric statistics comparison tests have been applied. Statistical analysis was carried out using OriginPro2020 SR1, MATLAB R2019b, and GraphPad Prism 6.07 & 9.

